# Reverse GWAS: Using Genetics to Identify and Model Phenotypic Subtypes

**DOI:** 10.1101/446492

**Authors:** Andy Dahl, Na Cai, Arthur Ko, Markku Laakso, Päivi Pajukanta, Jonathan Flint, Noah Zaitlen

## Abstract

Recent and classical work has revealed biologically and medically significant subtypes in complex diseases and traits. However, relevant subtypes are often unknown, unmeasured, or actively debated, making automatic statistical approaches to subtype definition particularly valuable. We propose reverse GWAS (RGWAS) to identify and validate subtypes using genetics and multiple traits: while GWAS seeks the genetic basis of a given trait, RGWAS seeks to define trait subtypes with distinct genetic bases. Unlike existing approaches relying on off-the-shelf clustering methods, RGWAS uses a bespoke decomposition, MFMR, to model covariates, binary traits, and population structure. We use extensive simulations to show these features can be crucial for power and calibration. We validate RGWAS in practice by recovering known stress subtypes in major depressive disorder. We then show the utility of RGWAS by identifying three novel subtypes of metabolic traits. We biologically validate these metabolic subtypes with SNP-level tests and a novel polygenic test: the former recover known metabolic GxE SNPs; the latter suggests genetic heterogeneity may explain substantial missing heritability. Crucially, statins, which are widely prescribed and theorized to increase diabetes risk, have opposing effects on blood glucose across metabolic subtypes, suggesting potential have potential translational value.

**Author summary:** Complex diseases depend on interactions between many known and unknown genetic and environmental factors. However, most studies aggregate these strata and test for associations on average across samples, though biological factors and medical interventions can have dramatically different effects on different people. Further, more-sophisticated models are often infeasible because relevant sources of heterogeneity are not generally known *a priori.* We introduce Reverse GWAS to simultaneously split samples into homogeneoues subtypes and to learn differences in genetic or treatment effects between subtypes. Unlike existing approaches to computational subtype identification using high-dimensional trait data, RGWAS accounts for covariates, binary disease traits and, especially, population structure; these features are each invaluable in extensive simulations. We validate RGWAS by recovering known genetic subtypes of major depression. We demonstrate RGWAS is practically useful in a metabolic study, finding three novel subtypes with both SNP- and polygenic-level heterogeneity. Importantly, RGWAS can uncover differential treatment response: for example, we show that statin, a common drug and potential type 2 diabetes risk factor, may have opposing subtype-specific effects on blood glucose.

## Introduction

Distinguishing subtypes can be essential for treatment, prognosis, and learning basic disease biology. For example, breast cancer has subtypes distinguished by tumor hormone receptor status that have different genetic risk variants, population structure, comorbidities, treatment responses and risks, and prognoses [1,2]. Many other common diseases have known, biologically distinct subtypes [3-7], often involving distinct tissues or biological pathways, including two diseases we study: depression [8] and type 2 diabetes (T2D) [9,10]. Genetically distinct subtypes can arise from gene-environment interactions [11-13]; gene-gene interactions [14,15]; or disease misclassification, which is well documented but usually ignored [16].

We aim to learn and validate genetic subtypes in a two-step approach we call reverse GWAS (RGWAS). First, we infer subtypes by clustering multiple traits with a new finite mixture of regressions method we designed specifically for large, multi-trait GWAS datasets (MFMR). The core assumption of MFMR is that the subtypes differ in distribution for many traits. Second, we assess the causal biological distinction between the subtypes by testing for SNP-and polygenic-level effect heterogeneity. We also test heterogeneity for non-genetic covariates, like medical interventions, which can distinguish the subtypes pragmatically [17-20]. These tests can be more powerful because genetic effects are usually small.

Unlike recent approaches to uncover subtypes [21-30], the first RGWAS step corrects for population structure, handles binary traits, and scales to tens of thousands of samples. Distinct from recent, complementary tests for genetic heterogeneity [22,31-34], the second RGWAS step offers p-values, polygenic tests, and covariate adjustment.

After we use genetics to validate inferred subtypes, we use the subtypes in turn to increase explained heritability, uncover genetic and treatment heterogeneity, and increase GWAS power. These applications resemble methods that infer heterogeneity from, for example, population structure [35] or technical artifacts [36,37].

We first introduce the two-step RGWAS approach. We then evaluate extensive simulations and find that RGWAS, but not several other previous and novel approaches, is calibrated and powerful. We then validate RGWAS in real data by recovering known SNP interactions with stress in major depression. Finally, we use RGWAS in a metabolic cohort and uncover three novel subtypes with polygenic, SNP, and treatment effect heterogeneity: subtypes increase explained heritability by ∼65%; identify 4 subtype-specific SNP effects, including 3 with known metabolic interactions; and find that statin, a widely prescribed drug that may increase diabetes risk [38-40], has opposing effects on blood glucose across subtypes.

## Results

### Reverse GWAS is calibrated and powerful in simulations

We simulate from the full MFMR model to assess RGWAS (Methods, Supplementary Section 3). We add noise that is correlated across traits; large main subtype effects; and a covariate matrix G containing null, homogeneous, and heterogeneous SNPs. We use 27 quantitative traits and 3 binary traits and simulate K = 2 subtypes (or K =1).

We test several other methods to find subtypes in step 1 (Methods). First, we use Gaussian Mixture Models (GMM) to represent covariate-unaware methods, e.g. k-means [27,30] and TDA [21,25,26]. Second, we use a new Canonical Correlation Analysis (CCA) approach that defines the subtype vector *z* as the top phenotypic CC. Third, we use the true *z* to show the best-case scenario with perfect subtyping (Oracle).

We then use *z* in the step 2 heterogeneity test, defined by (3), conditioning on the main effects of *z* (Figure 1). We aggregate 1,000 simulated datasets and, per dataset, the quantitative traits and SNPs in each category. Binary traits give similar results (Supplementary Figure 2).

**Fig 1.**
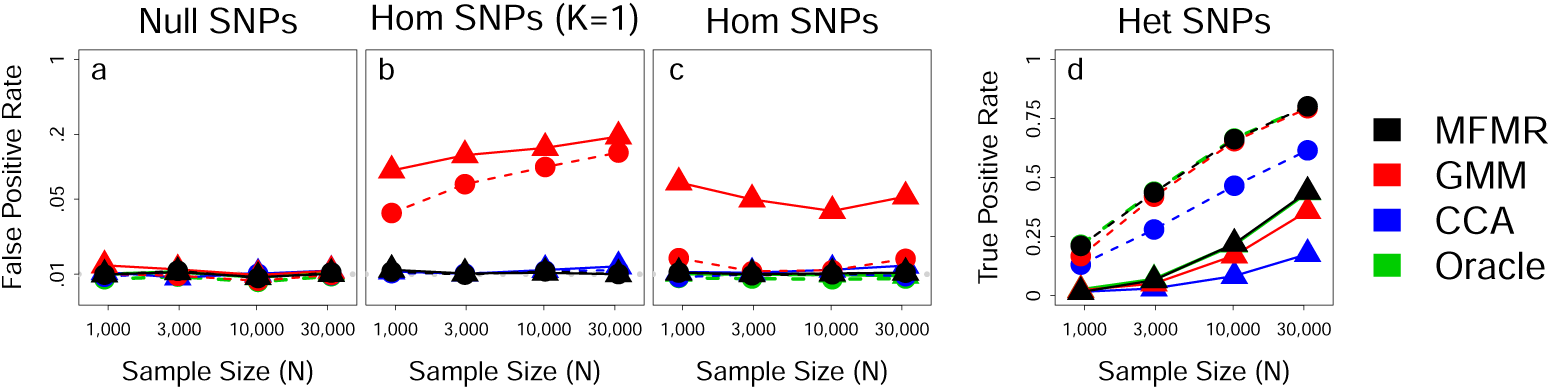
SNP heterogeneity tests at nominal. *p* =.01. SNPs are either null (*a*); homogeneous (hom, c); or truly heterogeneous (het, d); we also test hom SNPs in simulations with no subtypes (b). Hom SNPs explain 4% of variance and het SNPs explain .4% (triangles) or vice versa (circles).

RGWAS with MFMR is calibrated and almost perfectly obtains the oracle subtypes (Supplementary Figure 3) and power (Figure 1d). Crucially, it remains calibrated even when *K* = 1 (Figure 1b), so RGWAS discoveries validate the existence of subtypes. Further, when *K* > 2 subtypes were simulated, MFMR with fixed *K* = 2 lost power but remained calibrated (Supplementary Figure 4).

Conversely, GMM is miscalibrated by an order of magnitude when *K*=1, making it unreliable for subtype validation (Figure 1b). GMM cannot distinguish covariate from subtype, and it is inflated when homogenoues effects are stronger than heterogeneous (Figure 1c, Supplementary Figure 4).

CCA has low power but seems calibrated and, sometimes, to outperform the oracle (Supplementary Figure 4). But this is a Pyrrhic victory: by smoothing over traits, CCA causes bias, which adds signal for heterogeneous traits but inflates FPR (Supplementary Figure 5 and Section 4).

We also tried the top phenotypic PC for *z*, which performed like a lower-power CCA (Supplementary Figure 1). The top genetic PC [24], instead, had very low power.

We next simulated “Case/Control” data by ascertaining a binary trait to have 50% in-sample prevalence (Supplementary Figure 2). Little changed qualitatively, though all methods, even the oracle, had slight FPR inflation for large N because ascertainment violates the assumed model [41].

Intuitively, subtypes are clusters of covariate-adjusted traits. Variables that confound subtype structure, then, should included as covariates, while variables that may have different distributions between subtypes should be included as traits. (Covariates are ignored by methods like GMM.) Nonetheless, this distinction can be unclear in practice, which we test by treating a covariate like a trait (or vice versa). MFMR remained calibrated, unlike GMM (Supplementary Figure 6).

Finally, we simulated an even mixture of two populations and 10,000 SNPs from a Balding-Nichols model with F_st_ = .1 (Supplementary Section 3.2). We repeated our simulations using 12 simulated SNPs for G and adding population main effects of varying strength. For MFMR and the oracle, we condition on three genetic PCs and their interactions with *z* in step 2. MFMR remains calibrated and powerful while CCA and GMM suffer substantial FPR inflation, even for completely null SNPs (Supplementary Figure 2). Including PCs in step 2 after using GMM in step 1 can partially reduce inflation, but only when subtypes are truly present.

### RGWAS recovers known CONVERGE subtypes

Only one GWAS of clinical depression has reported replicated associations with major depression (MD) [42]. A possible explanation for this lack of GWAS hits is genetic, environmental, and/or diagnostic heterogeneity, suggesting RGWAS may be useful.

The CONVERGE study recruited and deeply phenotyped Han Chinese women with recurrent MD and matched controls. Cases were carefully ascertained to minimize environmental heterogeneity, comparatively amplifying biological heterogeneity. We used N = 9,303 samples measured on 31 binary and 10 quantitative traits. We conditioned on an intercept, age, and ten genetic PCs as heterogeneous covariates. We jointly imputed the covariates and traits (Methods).

The inferred subtypes with *K* = 2 are summarized in Figure 2. As expected, the subtypes distinguish lifetime adversity: the aggregate measure “Stress” is split [43].

**Fig 2.**
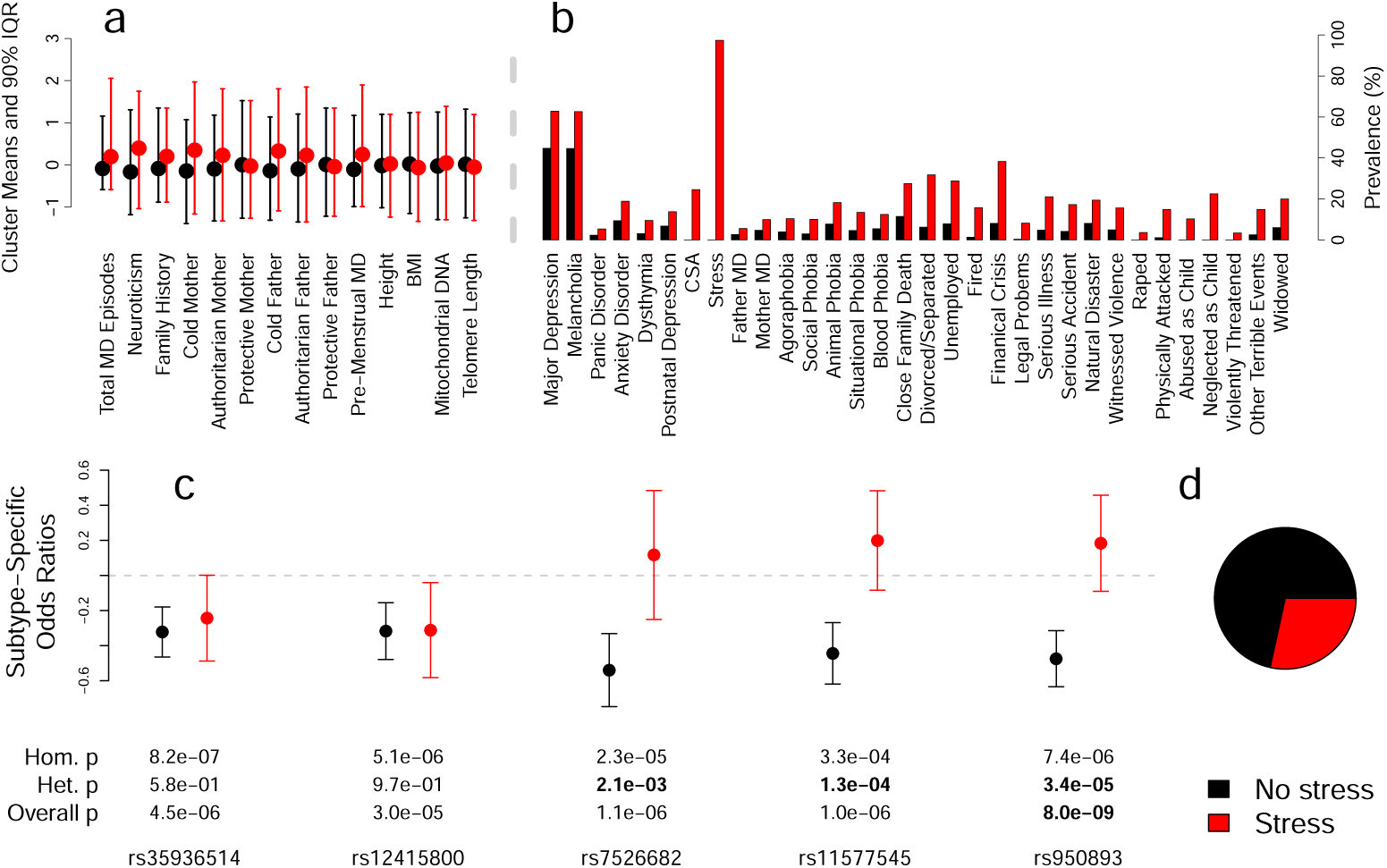
Genetic heterogeneity in the CONVERGE major depression dataset. Quantatitative trait 90% inter-quantile ranges (a) and binary trait prevalences (b) are shown for each subtype. (c) Per-subtypes odds ratios (±2 s.e.) for two SNPs discovered by (homogeneous) GWAS [42] (left) and three SNPs discovered using known subtypes [43] (right). (d) Subtype sizes.

We tested five SNPs for effect heterogeneity across subtypes (Figure 2c). The first two (rs35936514 and rs12415800) were discovered in the initial GWAS [42] and we use as negative controls for heterogeneity. For positive controls, we use three SNPs (rs7526682, rs11577545, and rs950893) previously found to interact with “Stress” [43]. As expected, the homogeneous SNPs are nearly genome-wide significant, and RGWAS successfully assigns all five SNPs. The heterogeneous SNPs show only modest homogeneous signal because they have essentially no effect in the “Stress” subtype.

We chose *K* = 2 using prior knowledge that MD can be split by (binary) stress. We assessed this empirically by evaluating the MFMR likelihood on held-out data, which supported *K* > 1 subtypes (Supplementary Figure 7). *K* = 3 creates an MD-only subtype, so we do not pursue *K* ≥ 3.

### New metabolic subtypes with genetic and pragmatic significance

We next applied RGWAS to metabolic traits measured in METSIM [44]. By combining genetic, environmental, metabolomic, and disease measurements, METSIM allows studying the pathway from risk factors to metabolic consequences to altered disease risk.

We studied 6,248 unrelated Finnish men. We used three binary traits: 854 samples had T2D, 3,524 had pre-diabetes (preT2D), and 541 had coronary heart disease (CHD); we excluded 15 samples with T1D. We used 13 quantitative traits, including 6 PCs of 228 nuclear magnetic resonance (NMR) metabolite measurements (capturing 77% of variance). As covariates, we used three genetic PCs, age, age^2^, and smoking, alcohol, statin, diuretic, and beta-blocker use.

We study K = 3 to compromise between parsimony and the cross-validated log-likelihood, which broadly supports 1 <K ≤ 8 (Supplementary Figure 7). To test robustness to perturbations, we used five-fold cross-validation. We found that 93% of originally co-clustered pairs (i.e. same most likely subtype) remained together, showing that people from the same population can be accurately assigned to existing subtypes.

The three inferred metabolic subtypes are summarized in Figure 3. They primarily distinguish the metabolomic PCs, which are aggregates of 228 NMR traits. To elucidate the subtypes, we fit logistic regressions on the raw NMR traits, conditional on statin, and studied those with nominal *p* <.01 (despite [27,29], these *p*-values are not calibrated). We first compared the large blue group to the combined orange and green groups, which suggested the blue group had less esterified cholesterol in small HDL and higher histidine and relative amounts of omega-3 fatty acid. Next, comparing orange to green indicated orange had more free but less esterified cholesterol, especially in large LDL, and that orange has more polyunsaturated fats and phenylalanine.

**Fig 3.**
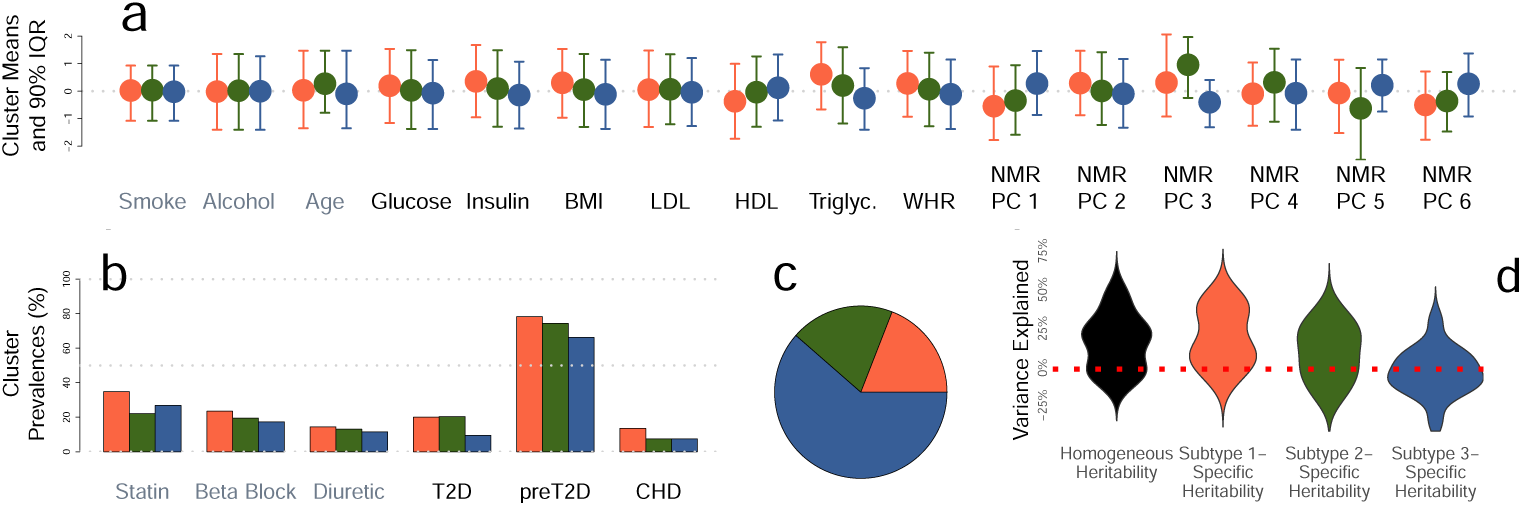
Three inferred metabolic subtypes in METSIM. (a) Quantitative and (b) binary distributions for covariates (grey labels, left) and traits (black lables, right). (c) Subtype sizes. (d) Across-trait distribution of ordinary (black) and additional subtype-specific heritabilities (colors) from Free GxEMM.

### Genetic metabolic heterogeneity

To test for polygenic subtype heterogeneity, we first used IID GxEMM and found significant subtype-specific heritability for 12/16 traits (p <.05/16, Figure 4),biologically distinguishing the subtypes. Further, this analysis increases average heritability estimates from 22.3% to 36.7% (Supplementary Figure 8), showing unknow subtypes can mask substantial heritability. As fixed effects, we used subtype main effects, age, age^2^, and three genetic PCs; for CHD, LDL, HDL, TG, and the NMR PCs,we also included statin. Variance explained is calculated after residualizing fixed effects [45]. The richer Free GxEMM fits significantly better than IID for 11/16 traits(p < .05/16, Supplementary Figure 8) and suggests, on average across traits, that orange and green subtypes have higher heritability than blue, which has roughly zero specific heritability (Figure 3d).

**Fig 4.**
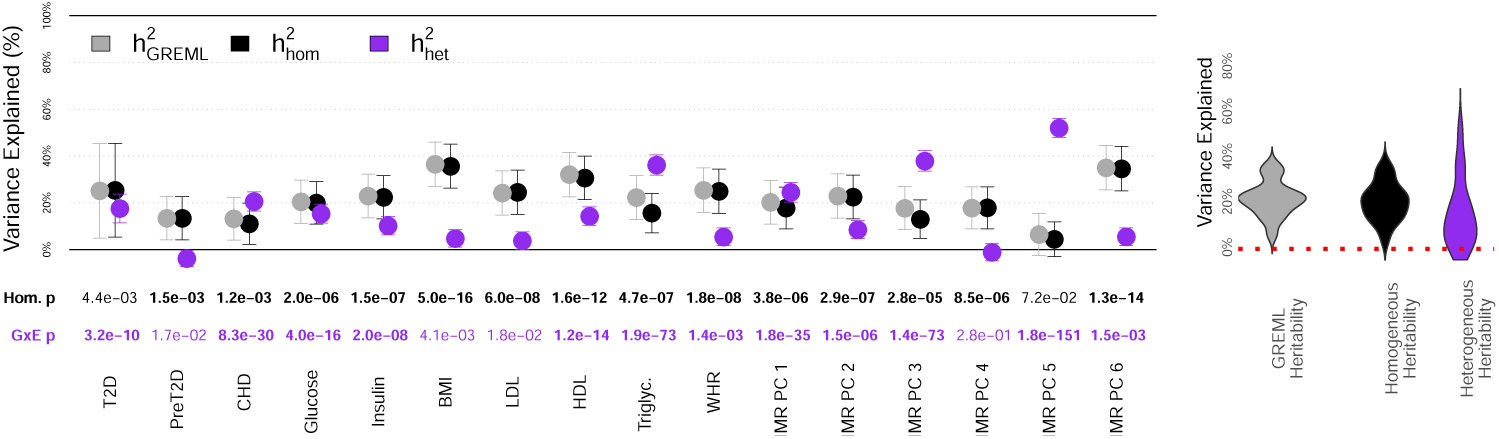
Polyge nic het ero geneit y in the inferred metabolic subtypes. Left:Point estimates ± 2 s.e. Right: Across-trait estimate distribution.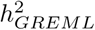 is the standard heritability estimate [46]. 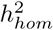and 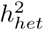 are the heritability estimates from IID GxEMM. Variance explained is on the observed scale for binary traits. The ‘Hom.’ *p*-value tests 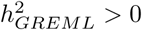;the GxE p-value tests 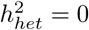.

With ∽6,000 samples, we do not have power for genome-wide heterogeneity tests. Instead, we test known metabolic GWAS SNPs-68 from T2D and 13 from CHD (Methods). We found four heterogeneous SNP-trait associations at *p* = .05/81. The orange and green effect estimates had opposite sign for 2/4, and all blue estimates were near zero. Together with the Free GxEMM results, this suggests that the blue group is a type of baseline and that partially overlapping biological pathways are specifically activated in the smaller groups.

These SNPs have several known metabolic interactions. rs10401969 is a splice variant for *SUGP1* that affects downstream splicing in the gene targeted by statins, *HMGCR* [47]; it also interacts with an APOE SNP on fenofibrate response [48]. rs7138803 interacts with exercise for obesity [49] and features in an obesity score interacting with diet [50]. rs780094 interacts with another SNP for fasting glucose [51], suggestively interacts with diet [52], broadly affects lipid levels, and is one of three SNPs in a risk score interacting with postprandial and post-fenofibrate cholesterol [53].

We next performed a genome-wide scan with the global, K df test (GxEWAS). This cannot establish SNP heterogeneity, but it can increase power over GWAS when heterogeneity exists. GxEWAS and GWAS give largely consistent results (Table 1, Supplementary Figure 9), as expected because the homogeneous and global tests are not independent. Nonetheless, GxEWAS is a valuable complement to GWAS as it discovers 10 additional loci (though GxEWAS misses 19/60 GWAS loci).

**Table 1.** Number of genome-wide significant loci for GWAS and GxEWAS. Bold indicates a locus not discovered by the other method (*r*^2^ < .2). No loci were found for CHD, insulin, or WHR. Both approaches found a single preT2D locus. We excluded NMR PC 5 because of GxEWAS inflation.

|  | Gluc. | BMI | LDL | HDL | TG | PC 1 | PC 2 | PC 3 | PC 4 | PC 6 |
| --- | --- | --- | --- | --- | --- | --- | --- | --- | --- | --- |
| GWAS | **3** | 0 | **7** | **7** | **8** | **6** | **4** | 0 | **5** | **19** |
| GxEWAS | **4** | **1** | 6 | **7** | **4** | **2** | **5** | **1** | 4 | 16 |
| Both | 2 | 0 | 6 | 5 | 3 | 1 | 3 | 0 | 4 | 16 |

To mimic prior approaches, we repeated the GxEWAS with covariate-unaware tests and GMM subtypes. Genome-wide QQ-plots were highly inflated for *K* ∊ {2, 3, 4} (Supplementary Table 1). Notably, *λ*_GC_ was even inflated for the binary traits, which we excluded from GMM so it would converge. This inflation can easily be mistaken for strong, ubiquitous signal when evaluating only candidate SNPs. The RGWAS *λ*_GC_ were comparatively modest, with a maximum of 1.34 (after excluding NMR PC 5, with *Xqc* = 1.83). Despite modest inflation for some traits, RGWAS substantially outperforms existing methods. Similar conclusions hold for the global, K df test.

### Pragmatic metabolic heterogeneity

We tested for statin effect heterogeneity to assess the pragmatic value of the metabolic subtypes. Using our test for large-effect covariates, only glucose had significant statin heterogeneity at *p* = 0.05/16 (*p* = 1.0 × 10^-4^); this was even clearer conditional on T2D *(p* = 2.5 × 10^-6^). There is no obvious FPR inflation as statin only has one other significantly heterogeneous effect across other traits and K ∊ {3,4, 5}. This is consistent with statin interactions with age [38] and genetically predicted LDL [40] on T2D, and also fenofibrate’s interaction with lipid levels on cardiovascular risk [54]. By contrast, large meta-analyses did not find inter-study statin heterogeneity [38,39].

To provide calibrated *p*-values, these analyses used the large-effect RGWAS test where MFMR treats statin as homogeneous. This test demonstrates statin effect heterogeneity in METSIM, which is further supported by tests with K = 4 and K = 5 (T2D-adjusted heterogeneity p =1.2 × 10^-5^ and 7.6 × 10^-4^, respectively). The *p*-value is insignificant for K = 2, indicating insufficient resolution.

We next tested statin heterogeneity with our primary metabolic subtypes (derived treating statin as heterogeneoues inside MFMR) with a heterogeneous linear regression on glucose, conditioning on our standard covariates and T2D (Figure 5b). The results indicate statin increases blood sugar in most people-consistent with [38,39]-but also that it may decrease glucose in the smaller, higher-risk orange and green groups.

**Fig 5.**
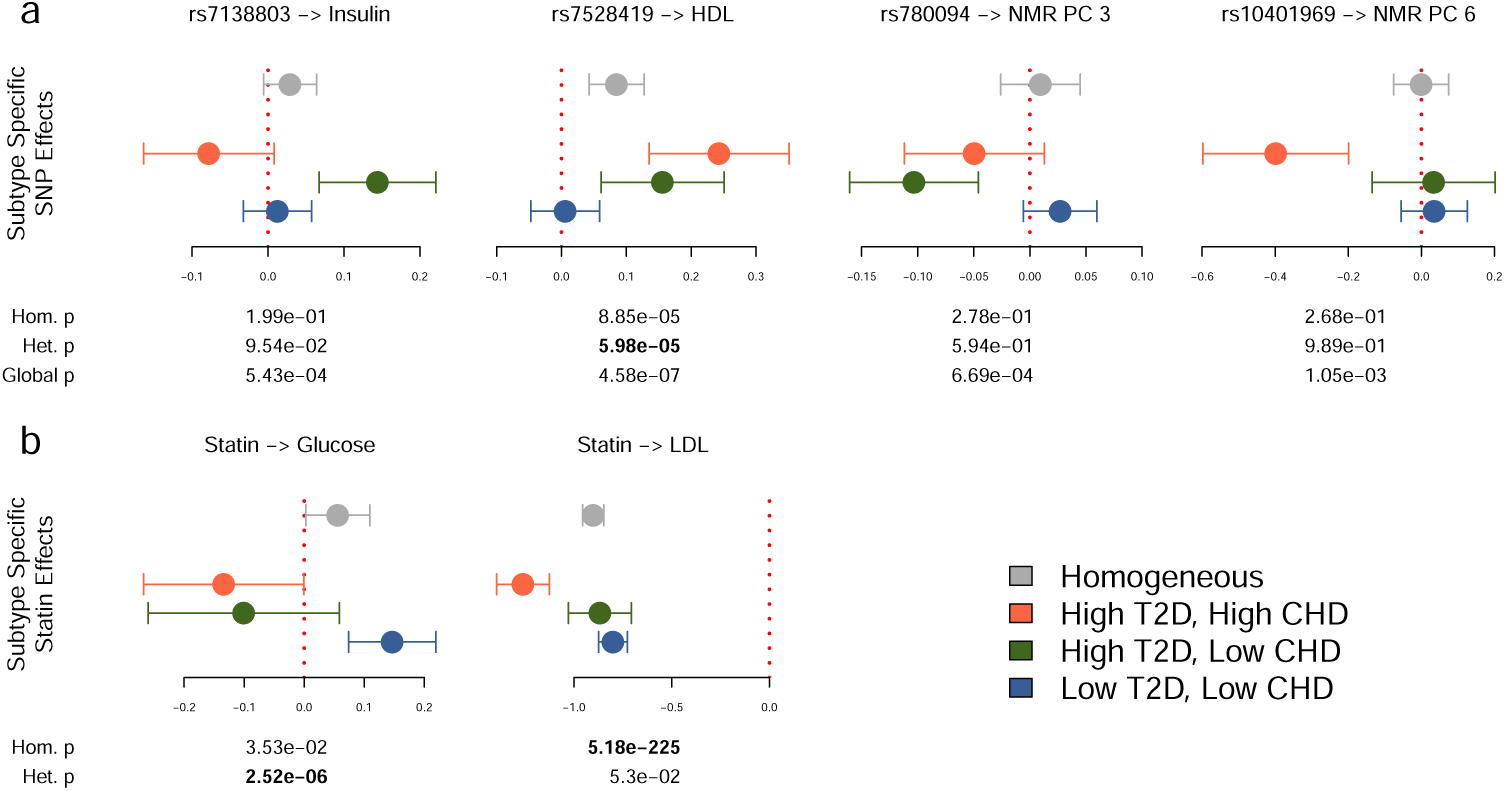
Significant metabolic subtypes effect heterogeneity. Subtype-specific effect estimates shown ± 2 s.e. (a) SNPs found with MFMRX test on 81 metabolic SNPs (p = .05/81). (b) Heterogeneous statin effect on blood glucose found with MFMR test on 16 traits (p = .05/16); statin LDL effect seems, misleadlingly, heterogeneous. For interpretation, we show estimates using on our main metabolic subtypes.

Since METSIM measured two time points, we tested the predictive value of our baseline subtypes for conversion from preT2D to T2D. We fit logistic regression on time 2 T2D status for the 1,924 baseline prediabetics. Subtypes significantly correlated with T2D conversion (*p* = 0.003), with orange and green converting less than blue. This remained true when conditioning on our standard covariates (*p* = .031). This also shows the subtypes persist over time, unlike prior, directly age-dependent T2D subtypes [55].

### Subtype-Specific Tissue Heritability Enrichment

Some traits, including metabolic diseases [15,56], have disparate genetic effects acting through distinct cell types, tissues, or biological processes. Subtypes that differentiate biological modes of action at this systems-level would be more easily interpretable and useful for basic research and precision treatment.

We used LDSC-SEG to partition subtype-specific heritability across GTEx tissues for several traits (Supplementary Section 5) [57]. We use a merely suggestive *p* = .01 threshold rather than attempting to account for testing multiple tissues and traits. First, homogeneous subtype meta-analysis found pancreas enrichment for NMR PC 5, LDL, insulin, and triglycerides (Supplementary Figure 10). Second, heterogeneous meta-analysis found five cell type-trait pairs with subtype-specific enrichments, including transformed fibroblasts-NMR PC 5, in line with the complex roles of cholesterol in skin cells and NMR PC 5, and frontal cortex-blood glucose.

## Discussion

In a purely descriptive sense, inferring subtypes is easy: applying any clustering algorithm to any data produces subgroups. But existing methods cannot go beyond such descriptions because they are liable to downstream FPR inflation. By contrast, RGWAS is calibrated in simulation, recovers known MD subtypes, and produces biologically and pragmatically validated metabolic subtypes. RGWAS handles covariates, mixed binary and quantitative traits, residual trait correlations, and is implemented in the simple, free **rgwas**R package, available with a vignette at https://github.com/andywdahl/rgwas.

There are several limitations to RGWAS. First, like other two step methods, RGWAS fails to propagate first-step uncertainty. Similarly, although we do not imagine there is a “true” K, more can always be done to better choose K. Also, while we have tested a variety of simple decompositions to learn subtypes, others may perform better, especially where domain-specific tools exist. In particular, MFMR is conceptually similar to a matrix factorization/depth-two linear network, suggesting inner layers of appropriate neural networks may define useful subtypes.

There are also specific limitations to our inferred stress subtypes in CONVERGE. First, our stress measurements were retrospective and self-reported, meaning our stress traits could be biased by MD status. Second, our analysis was not entirely without domain supervision because we included the aggregate trait “Stress” that was previously manually constructed [43]. Nonetheless, RGWAS identified the key trait amongst dozens, and our METSIM analysis demonstrates that RGWAS can be useful without any domain guidance.

MFMR is only a first step toward genetic subtyping, and there are many possible extensions. Sparsifying penalties can be incorporated by replacing CM steps with calls to third-party software and could extend MFMR to higher-dimensional traits and covariates. A random-effect version of MFMR could improve power to detect polygenic subtypes, though computational issues are non-trivial. MFMR could also be adapted to count data, zero-inflation, higher-order arrays, or missing data. Theoretically, it would be interesting to let subtypes vary between traits, which MFMR can capture only with large K. Instead of an i.i.d. prior on *z*, MFMR could model *z* with a multinomial logistic regression to estimate, test, and correct for effects on *z*, which can be directly interesting [58] and can clarify the interpretation of heterogeneity [59]. Or, instead, we could use a continuous prior on *z* with a factor analysis model [22]. Finally, MFMR could be applied only within diseased individuals to directly define subtypes of disease; however, this requires fundamentally different step 2 tests, and subtype heterogeneity is not generally disease relevant [31].

Our polygenic approach to subtype validation with GxEMM provides a much needed power advantage over SNP-level heterogeneity tests at the cost of resolution; conceptually, polygenic risk score tests lie between [33,60]. But SNP-level precision is not needed to meet our criterion for biologically meaningful subtypes, making GxEMM invaluable for subtype validation. Nonetheless, its assumed linear model can confuse non-linear effects for heterogeneity. Similar issues arise in generalized linear models, as the existence of effect heterogeneity depends on link function. In particular, tests for differential disease heritability give different results on the liability and observed scales [61,62], reducing confidence in our GxEMM results for T2D and CHD. Particular forms of non-linearity, e.g. ascertained binary traits, can be accommodated under genetic homogeneity [63-68], which may be extensible to heterogeneity.

Although we focused on fixed- and random-effect interaction tests to establish heterogeneity between subtypes in step 2, it may also be useful to apply recent, complementary heterogeneity tests. For example, Subtest could be used to assess differences between K = 2 disease-only subtypes [31]. For large K, on the other hand, StructLMM is a natural complement to GxEMM: the latter is more powerful because it uses genome-wide information and a richer GxE model, but the former has SNP-level resolution and scales to dramatically larger N and K. Similarly, large-K subtypes could be post-processed with hierarchical clustering and tested with TreeWAS [33]. Broadly, any heterogeneity test can be used in the second step, as we demonstrated with our LDSC-SEG application in METSIM.

While we did assess tissue-specificity in our metabolic clusters in step 2, we did not actively encourage subtypes to be differentiate tissues in step 1. In the future, we will do this by incorporating tissue specific genetic risk scores as heterogeneous covariates inside MFMR, which will allow the algorithm to prioritize tissue-specific subtypes.

Finally, as MFMR seeks clusters that are unaffected by confounders like population structure, age or sex, it may be useful for clustering in settings where protecting certain information is important for privacy or fairness [69]. In this sense, MFMR is to GMM roughly as AC-PCA [70] or contrastive PCA [71] are to ordinary PCA.

## Methods

### Ethics Statement

#### CONVERGE

The study protocol was approved centrally by the Ethical Review Board of Oxford University (Oxford Tropical Research Ethics Committee) and the ethics committees of all participating hospitals in China. All participants provided written informed consent.

#### METSIM

The Ethics Committee of the University of Eastern Finland and Kuopio University Hospital approved the METSIM study, and this study was conducted in accordance with the Helsinki Declaration. All participants provided written informed consent.

#### RGWAS step 1: clustering with MFMR to find subtypes

We derive a novel clustering algorithm, multitrait finite mixture of regressions (MFMR), beginning from the standard regression model for interaction. Assuming a quantitative trait *y*, covariates X, discrete subtypes z, and a focal covariate *g* putatively interacting with *z*, the model is:

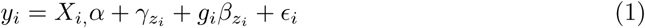

X_*i*_, is a vector of Q control covariates, like genetic PCs or sex, with homogeneous effect sizes *a. zi* ∊ {1,…, K} is a K-level factor specifying the subtype for individual *i*, and *γ_k_*are its main effects.β is the vector of subtype-specific *g* effects. We say *g* is homogeneous if βι =…= β_κ_; otherwise, *g* is heterogeneous. We assume ∊ is i.i.d. Gaussian with mean zero.

MFMR generalizes (1) in several complementary directions. First, we allow a matrix of heterogeneous covariates-G instead of *g.* Second, we learn the subtypes (*z*) instead of assuming they are known (giving a Finite Mixture of Regressions, FMR) by assuming *z_i_* are i.i.d. Categorical:

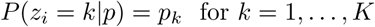

Third, we generalize *y* a matrix Y of Multiple traits (MFMR), which adds power for subtypes that affect the distribution of many traits. This power is crucial in practice because genetic interactions are often weak.

Finally, we model binary traits with probit link functions to mitigate the spurious local modes that plague methods like *k*-means. For example, this issue led others to discard roughly half their data *post hoc* [30]. This model is computationally prohibitive even for modest B, which we address with a novel conditional independence assumption. This induces constraints in our optimization which we solve with block matrix identities (Supplementary Section 2.3).

We fit MFMR with an Expectation Conditional-Maximization (ECM) algorithm. Our ECM generalizes standard EM for Gaussian Mixture Models. Both iterate between *z* updates in E-steps and parameter updates (e.g. α and β) in (C)M steps.

When fitting MFMR in step 1, a covariate that will be tested for heterogeneity in step 2 can either be ignored (MFMRX), included in X (MFMR, our default), or included in G (MFMR+). In Gaussian mixture models, covariates can only be ignored (GMM) or added as traits (GMM+) [30]. MFMR+ and GMM+ overfit in simulations, causing false positive rate (FPR) inflation (Supplementary Figure 1). Conversely, MFMRX and GMM underfit homogeneous covariates, which also inflates FPR (Figure 1). MFMR strikes a balance: the homogeneous effect is adjusted but subtypes are not tuned to the heterogeneous effect. This resembles a score test as the alternate is tested by fitting only the null. However, small-effect covariates, like SNPs, can be safely ignored [72], enabling genome-wide testing with MFMRX.

We note that MFMR generalizes several well-known models. If binary traits and X are excluded and the covariates G are reduced to an intercept, MFMR becomes GMM. When P =1 and *z* is known, MFMR becomes a standard gene-environment interaction (GxE) model with discrete environments/subtypes. Finally, if P =1 and *β_k_* = β_0_ for all *k*, MFMR reduces to linear/probit regression.

#### RGWAS step 2: calibrated tests to validate subtypes

For simplicity, we assume there are K = 2 subtypes and just one interacting covariate, *g.*These assumptions mean the output from step 1 is just a vector *z*, where *z_i_* is the subtype 1 probability for sample i, and that the interaction model takes a simple form:

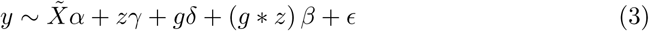

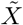 collects all background covariates, like genetic PCs, unlike existing subtype validation tests that largely ignore population structure [23,25,28,30]. * is element-wise multiplication, but it can be generalized to allow K > 2 and a matrix G instead of a single covariate g.

We consider three tests for g: the homogeneity test for δ ≠ 0 given β ≠ 0; the heterogeneity test for β ≠ 0 with free δ; and the global test for δ, β ≠ 0 [73]. The homogeneity test has 1 degree of freedom (df), the heterogeneity test has K — 1 df, and the global test has K df. We focus on the heterogeneity test, which establishes that g has differential effects across subtypes and thus that the subtypes differ in causal biology (if g is genetic) or pragmatically (e.g. if g is a treatment). We assume ∊ is i.i.d and test with linear or logistic regression.

We also use a polygenic version of (3), GxEMM [62], to jointly model and test β across all SNPs with random effects. GxEMM uses a predefined inter-sample genetic similarity matrix [45,74] to partition phenotypic heritability into a component shared between subtypes 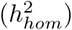 and subtype-specific components. GxEMM is useful for genetic subtyping because the test 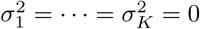 is a powerful way to demonstrate that subtypes have partially distinct genetic bases when sample size is too low to discover individual subtype-specific SNP effects. We fit both Free GxEMM, which learns specific heritabilities in each subtype 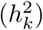, and IID GxEMM, which assumes subtypes have equal heritability 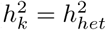for all *k*).

### Other approaches to infer subtypes

We develop a novel subtyping approach by applying CCA to G and the joint binary and quantitative phenotype matrix (Y^b^: Y), both centered and scaled, and taking *z* to be the top phenotypic CC. CCA (and phenotypic PCA) defines *z* as a linear trait combination, implying its heterogeneity tests cannot be trait-specific (Supplementary Section 4). Nonetheless, sparse estimators can resolve this problem in theory, and CCA is computationally efficient (Supplementary Figure 3).

We also tested GMM, which models samples as draws from one of K multivariate Gaussians. We fit GMM to the quantitative traits with a standard EM algorithm [75]. We consider GMM similar, in the sense of covariate-unawareness, to k-means, which struggles even more with binary traits, and TDA, a proprietary package.

Most similar to MFMR, LIMMI aims to identify GxE with unknown E in gene expression [22]. Beyond many technical differences, LIMMI and MFMR are built for disjoint scenarios: MFMR only fits tens of traits, but LIMMI only fits hundreds of samples, preventing its use in our setting.

### METSIM dataset

We selected metabolically relevant SNPs by taking published GWAS SNPs for T2D or CHD. We used the 153 T2D SNPs in Table 1 of [76] as known T2D SNPs. We had genotyped 86 of these SNPs, which we reduced further to 68 roughly independent SNPs (*r*^2^<.1). We used the 65 CHD SNPs in Supplementary Table 2 of [77] as known CHD SNPs, 13 of which we genotyped (all *r*^2^< .1). We filtered the original 10,070 person dataset so all pairwise kinships were below 0.05, as in [45].

### Phenotype imputation

We imputed missing data before running MFMR in CONVERGE. We jointly imputed covariates and traits with a sample-wise i.i.d. Gaussian model (MVN-impute from [78]). We thresholded imputed entries in Y^*b*^ to {0,1} in order to retain the downstream logistic regression framework. By contrast, discarding samples with any missing data reduces sample size by roughly half and the known positive SNP interactions were no longer recovered.

We imputed METSIM similarly, including all 228 NMR traits at the imputation step. We used softImpute to accommodate the wide matrix [79].

## Acknowledgments

We thank members of Zaitlen lab for helpful discussions and the individuals who participated in the CONVERGE and METSIM studies. This work was funded by National Institutes of Health (NIH) grants 1U01HG009080-01, 5K25HL121295-03, 1R03DE025665-01A1, HL-095056, HL-28481, and U01 DK105561. N. Cai was supported by the EBI-Sanger Postdoctoral Fellowship. A. Ko was supported by the NIH grant F31HL127921.

## Supporting information

S1 File. Supplementary Note. Full description of MFMR model and EM algorithm. Also includes proofs about PCA/CCA subtype estimators.

S1 Fig. Main Figure 1 with further subtyping methods.

S2 Fig. Alternate versions of Figure 1. Heterogeneity tests applied to simulated binary traits, with and without ascertainment, and after adding population structure.

S3 Fig. Simulation runtime and clustering accuracy summaries.

S4 Fig. Heterogeneity test power varying further simulation parameters.

S5 Fig. Simulations where SNP heterogeneity varies between traits.

S6 Fig. Simulations where covariates and traits are misplaced.

S7 Fig. Out-of-sample against *K* in CONVERGE and METSIM.

S8 Fig. Polygenic heterogeneity in the inferred metabolic subtypes.

Comparison of the IID and homogeneous heritability estimates.

S9 Fig. Comparison of the −log _10_(*p*)-values for GWAS and GxEWAS.

S10 Fig. Cell type-specific enrichment in METSIM metabolic subtypes.

S1 Tab. *λ_GC_* for GWAS, MFMR GxEWAS, and GMM GxEWAS.

